# Protein thermal stability does not correlate with cellular half-life: Global observations and a case study of tripeptidyl-peptidase 1

**DOI:** 10.1101/828509

**Authors:** Aaron M. Collier, Yuliya Nemtsova, Narendra Kuber, Whitney Banach-Petrosky, Anurag Modak, David E. Sleat, Vikas Nanda, Peter Lobel

## Abstract

Late-infantile neuronal ceroid lipofuscinosis (LINCL) is a neurodegenerative lysosomal storage disorder caused by mutations in the gene encoding the protease tripeptidyl-peptidase 1 (TPP1). Progression of LINCL can be slowed or halted by enzyme replacement therapy, where recombinant human TPP1 is administered to patients. In this study, we utilized protein engineering techniques to increase the stability of recombinant TPP1 with the rationale that this may lengthen its lysosomal half-life, potentially increasing the potency of the therapeutic protein. Utilizing multiple structure-based methods that have been shown to increase the stability of other proteins, we have generated and evaluated over 70 TPP1 variants. The most effective mutation, R465G, increased the melting temperature of TPP1 from 55.6°C to 64.4°C and increased its enzymatic half-life at 60°C from 5.4 min to 21.9 min. However, the intracellular half-life of R465G and all other variants tested in cultured LINCL-patient derived lymphoblasts was similar to that of WT TPP1. These results provide structure/function insights into TPP1 and indicate that improving *in vitro* thermal stability alone is insufficient to generate TPP1 variants with improved physiological stability. This conclusion is supported by a proteome-wide analysis that indicates that lysosomal proteins have higher melting temperatures but also higher turnover rates than proteins of other organelles. These results have implications for similar efforts where protein engineering approaches, which are frequently evaluated *in vitro*, may be considered for improving the physiological properties of proteins, particularly those that function in the lysosomal environment.

## Introduction

Late-infantile neuronal ceroid lipofuscinosis (LINCL) is a lysosomal storage disease (LSD), and is one of the most frequently occurring members of the family of Batten diseases ^1,2^. Symptoms typically manifest around 2-4 years of age and include blindness, ataxia, neuro-regression, and seizures that increase in severity as the child ages. Left untreated, patients with LINCL typically have a lifespan of 6-15 years before the disease proves fatal. LINCL is caused by autosomal recessive mutations in the *TPP1* gene (formally designated *CLN2*) that result in the loss or decreased activity of the encoded lysosomal protease ^3^ that was subsequently shown to be tripeptidyl-peptidase 1 (TPP1) ^4,5^.

TPP1 is a member of the sedolisin (MEROPS S53) family ^6^. These unusual serine peptidases have an overall fold similar to subtilisin, but instead of a canonical Asp-His-Ser catalytic triad they have an Asp-Glu-Ser triad that allows the active site serine to function in nucleophilic attack of substrate at acidic pH ^6–8^. *TPP1* encodes a 563 aa preprotein comprised of a 19 aa signal sequence, 152 aa pro-domain (aa 20-171), 24 aa linker region (aa 172-195), and a 368 aa (aa 196-563) catalytic domain ^3,7,8^. The crystal structure of the human proenzyme reveals a suboptimal catalytic triad geometry with the pro-piece linker partially blocking the substrate binding site. This arrangement is believed to prevent premature activation of the enzyme ^7–9^. When exposed to pH ≤ 4.0, TPP1 undergoes endoproteolytic cleavage at one of three cleavage sites located along the linker region resulting in an enzymatically active mature protein ^7,10,11^. This is likely due to a rearrangement of the catalytic triad at low pH. Mature TPP1 is then fully active with a pH optimum of 4.5 – 5.0 ^12^. TPP1 possess three internal disulfide bonds (C111-C122, C365-C526, and C522-C537) an octahedrally coordinated calcium-binding site, and five glycosylation sites (Asn 210, 222, 286, 313, and 443) ^7,8,13,14^. TPP1 glycans contain mannose-6-phosphate (M6P) ^3,13^. This modification is recognized by M6P receptors which guide intracellular trafficking of newly-synthesized TPP1 to the lysosome, as well as the endocytic uptake and delivery of extracellular TPP1 to the lysosome where the pro-form of TPP1 (proTPP1) becomes enzymatically active ^11^. TPP1 has two enzymatic functions: a primary tripeptidyl exopeptidase activity where TPP1 catalyzes the sequential release of tripeptides from the *N*-terminus of a broad range of substrates ^15,16^ and a weak endoproteolytic activity ^12,17^.

The only approved therapy for LINCL is enzyme replacement based upon administration of recombinant human TPP1 to the cerebrospinal fluid of patients, resulting in a stabilization of clinical parameters compared to untreated patients ^18^. The approved treatment regimen in the US is 300 mg of recombinant enzyme injected biweekly and this frequency of administration places a significant compliance burden on patients. In addition, extremely high costs are associated with the production of the large amounts of recombinant enzyme required for this first-generation treatment. One route to improve this therapy would be to engineer a TPP1 variant with an increased functional half-life in the lysosome. This would enhance the potency of the therapeutic and potentially increase the efficiency of the treatment by reducing the amount of recombinant TPP1 required per treatment, and/or increasing the interval between doses.

Previous studies involving proteins such as L-asparaginase, β-galactosidase, λ-repressor, Clostridium difficile toxin A-specific antibodies, and *Bacillus subtilis* lipase have shown that increasing a protein’s thermostability may also increase its resistance to protease digestion ^19–22^. We therefore hypothesized that increasing the thermostability of TPP1 would enable it to remain enzymatically active for longer in the protease rich environment of the lysosome. In this study, we utilized multiple structure and sequence-based design approaches to potentially increase the thermodynamic stability of TPP1, with the ultimate goal of engineering active variants with increased stability within the lysosome. While we did identify mutations that increased TPP1 stability *in vitro*, these did not translate to increased physiological half-life. These results have implications for similar efforts in which protein engineering approaches might be considered for improving the biological properties of other eukaryotic proteins, particularly those that functional in the complex environment of the lysosome.

## Materials and Methods

### Molecular Cloning

The TPP1 coding sequence was previously cloned into a pmCherry-N1 vector backbone (Clontech, CA) in place of the *pmCherry* coding sequence ^23^. Desired changes were introduced utilizing “round the horn site-directed mutagenesis” ^24^ with a DpnI restriction digest included to remove parental template. Primers were purchased from Integrated DNA Technologies Inc. and molecular biology enzymes were purchased from New England Biolabs. Mutations were verified by sequencing (Macrogen) and DNA for transfection was purified using the PureYield Plasmid MaxiPrep System (Promega).

### TPP1 Activity Assay

ProTPP1 (aa 20-563) samples in either PBS or conditioned ExpiCHO media were preactivated for two hrs at 37°C in buffer containing 0.1 M sodium formate / 0.15M sodium chloride / 0.1% Triton X-100, pH 3.5 ^12^. Samples were further diluted and TPP1 activity was measured using an endpoint assay with an Ala-Ala-Phe-7*amino-4-methylcoumarin* substrate in 0.1 M sodium acetate / 0.15M sodium chloride / 0.1% Triton X100, pH 4.5 as previously described ^25^. Activity of dilutions in the linear range were compared to a WT TPP1 standard of known concentration (generously provided by BioMarin Pharmaceutical) that was processed in parallel. In terms of overall experimental design, TPP1 variants were assumed to have a specific activity equal to that of WT TPP1 in the initial screen described above, while rigorous structural, enzymological and pharmacokinetic analyses would be conducted on any variants with apparently improved physiological properties.

### Production of TPP1 Variants

TPP1 constructs were expressed using the ExpiCHO system using reagents and protocols from the supplier (ThermoFisher Scientific). Briefly, serum-free medium adapted CHO-S cells were cultured in suspension in 125 ml Erlenmeyer cell culture flasks (Corning) at 37°C and 8% humidified CO_2_ in 35 ml of ExpiCHO expression medium and transiently transfected with plasmid DNA. The following day, the cultures were placed at 32°C and 5% CO_2_ and penicillin/streptomycin was added. After 12 days, the conditioned media was collected for subsequent analysis. Successful variant expression was determined by the observation of TPP1 activity in the media using the TPP1 activity assay.

### Purification of TPP1

To facilitate the purification of proTPP1, variants of interest were engineered to contain a C-terminal hexahistidine tag to enable purification from conditioned media using HisPur Cobalt Resin (ThermoFisher Scientific). Sample purity was verified by the observation of only a single band on an SDS-PAGE gel following purification.

### Cellular half-life assay

LINCL lymphoblast line GUS15820 was derived from a patient that was compound heterozygous for 1946A → G and 3556G → C splice-junction mutations ^26^. The identity of the LINCL lymphoblasts was verified previously by sequencing as well as TPP1 activity assay. Lymphoblasts were grown in suspension at 37°C and 5% CO_2_ in RPMI, 15% fetal bovine serum, and 1X Penicillin/Streptomycin in T25 flasks. LINCL lymphoblasts were grown to a density of 500,000 cells/mL in eight mLs of media. For a given set of experiments, two biological replicates of each variant were analyzed in parallel with two biological replicates of a WT TPP1 control. TPP1 (~26.4 μg) was added to a final concentration of 50 nM and cells were cultured for two days to allow for TPP1 uptake, then centrifuged at 200 X G for 10 min. Media was removed and the cell pellet was washed with 1X PBS and resuspended in fresh media lacking TPP1. Cell samples were collected in duplicate (technical replicates) on the initial day as well as each of the subsequent four days with each sample washed with 1X PBS, frozen on dry ice, and stored at −20°C. Fresh media was added during the collection phase to account for cell doubling and collection volumes were adjusted accordingly to ensure that the same proportion of cells was sampled at each time point. After day 5, cell samples were lysed and TPP1 activity measured ^11^. LINCL lymphoblasts without the addition of TPP1 were used to determine background. The technical replicates from all time points were used to calculate a half-life for each biological replicate. Each variant was then normalized to the biological replicate(s) of the WT TPP1 analyzed in parallel to account for day to day differences in experimental conditions. Experiments were performed utilizing at least two biological replicates for each TPP1 variant.

### Circular Dichroism

Purified TPP1 proenzyme (final concentration 0.5 mg/ml) was incubated overnight in 0.1 M sodium formate / 0.15M sodium chloride/ 0.1% reduced Triton X-100 pH 3.5 with dialysis against the same buffer (20kDa MWCO, ThermoFisher) to convert TPP1 to a mature/active form and to remove the cleaved pro-domain. Absence of residual pro-domain was verified by MALDI-TOF mass spectrometry. Circular dichroism (CD) experiments were then carried out using an AVIV Model 420SF CD Spectrometer (Aviv Biomedical, Lakewood, NJ). CD wavelength spectra were measured from 210-260 from 5°C - 95°C in 10°C intervals with 10 min equilibration time using 10 sec averaging. For thermal denaturation experiments, ellipticity was monitored at 222 nm from 4°C - 90°C in 2°C intervals with 2 min equilibration time and 10 sec averaging. Blank buffer subtraction was performed on each sample.

### Thermostability assays

Conditioned media containing proTPP1 was diluted to a concentration of 1.0 μg/mL with complete RPMI medium, 15% FBS, 1X penicillin/streptomycin. The samples were preactivated by being diluted 10-fold in 0.1 M sodium formate / 0.15M sodium chloride / 0.1% Triton X-100, pH 3.5 and incubated at 37°C for two hrs. Samples where then diluted 10-fold in 0.5 M sodium acetate / 0.15M sodium chloride / 0.1% Triton X-100, pH 5.0, incubated at 60°C for 0, 5, 15, 30, 45, and 60 min, and then analyzed by TPP1 activity assay. Experiments were performed in duplicate, with two technical replicates obtained from each sample at each time point. These technical replicates were then averaged to obtain the final value.

### Statistics

Unless otherwise noted, statistical analysis was performed using GraphPad Prism version 5.01. TPP1 half-life at 60°C and in the lysosome were calculated using a one-phase exponential decay model, setting the final plateau value to 0. K_uptake_ values were calculated using a one-site specific binding model. T_m_ values were determined following the fitting of the CD/activity data to the Hill equation. The results are expressed as mean ± SD.

## Results

Our overall approach was to use principles of protein design largely directed at increasing the thermodynamic stability of mature human TPP1, with the hypothesis that this would result in a variant with a longer enzymatic half-life within the protease rich environment of the lysosome. Constructs were initially expressed by transient transfection in ExpiCHO cells. Samples of conditioned media were acidified to convert the secreted proenzyme into the mature active form and the concentration of active variants was estimated by comparison with purified WT recombinant human TPP1. Expressed variants possessing enzymatic activity were further analyzed for stability of the active form at 60°C and/or for half-life within the lysosome following uptake of proenzyme by LINCL lymphoblasts.

We employed eight different protein engineering approaches, the majority of which utilized the previously determined crystal structure of proTPP1 (PDB ID: 3EDY) ^7^ to identify TPP1 variants with potentially increased lysosomal stability. Most approaches focused on improving the thermodynamic stability either by stabilizing the folded state or destabilizing the unfolded state of TPP1. The strategies employed were: (1) increasing the conformational stability of β-turn structures by alleviating steric hindrance; (2) introducing additional, internal disulfide bonds; (3) increasing structural homology with the thermoacidophilic peptidase kumamolisin; (4) improved hydrophobic packing of the protein core; (5) introducing additional metal ion binding sites; (6) introducing additional salt bridges; (7) alteration of the primary structure to include “stabilizing dipeptides;” and (8) introducing additional potential N-linked glycosylation sites. Using these strategies, we generated over 70 TPP1 variants for subsequent evaluation.

For *in vitro* half-life experiments, TPP1 variants were activated at pH 3.5 for two hours and then exposed to 60°C, a temperature slightly higher than the 56.2°C T_m_ of mature TPP1 (see below), with samples taken at timed intervals and analyzed by TPP1 activity assay (Figure 1A). WT TPP1 had a median half-life of 5.3 min at 60°C. Experiments to determine the functional half-life of TPP1 variants in the lysosome were performed in lymphoblasts derived from LINCL patients which lack endogenous TPP1 ^26^. Consistent with earlier studies in LINCL fibroblasts ^11^, uptake of TPP1 by LINCL lymphoblasts occurred in a concentration-dependent manner with M6P inhibiting the high affinity uptake phase (Figure 1B). This indicated both endocytosis by M6P receptors with a K_uptake_ of 0.8 nM and a low affinity, non-specific fluid phase endocytosis (Figure 1B). To maximize specific uptake for the lysosomal half-life experiments, LINCL lymphoblasts were incubated with 50 nM of each TPP1 variant, after which the TPP1 containing media was removed and replaced with fresh media. Cell samples were collected at timed intervals with extracts assayed for TPP1 activity. WT TPP1 had a median lysosomal half-life of 42 hours (Figure 1C). A compendium of all of the TPP1 variants characterized and their respective functional properties is provided in Table S1. Strategies used to guide the design of each variant and the effect of the respective mutations on stability is described below.

**Figure 1.**
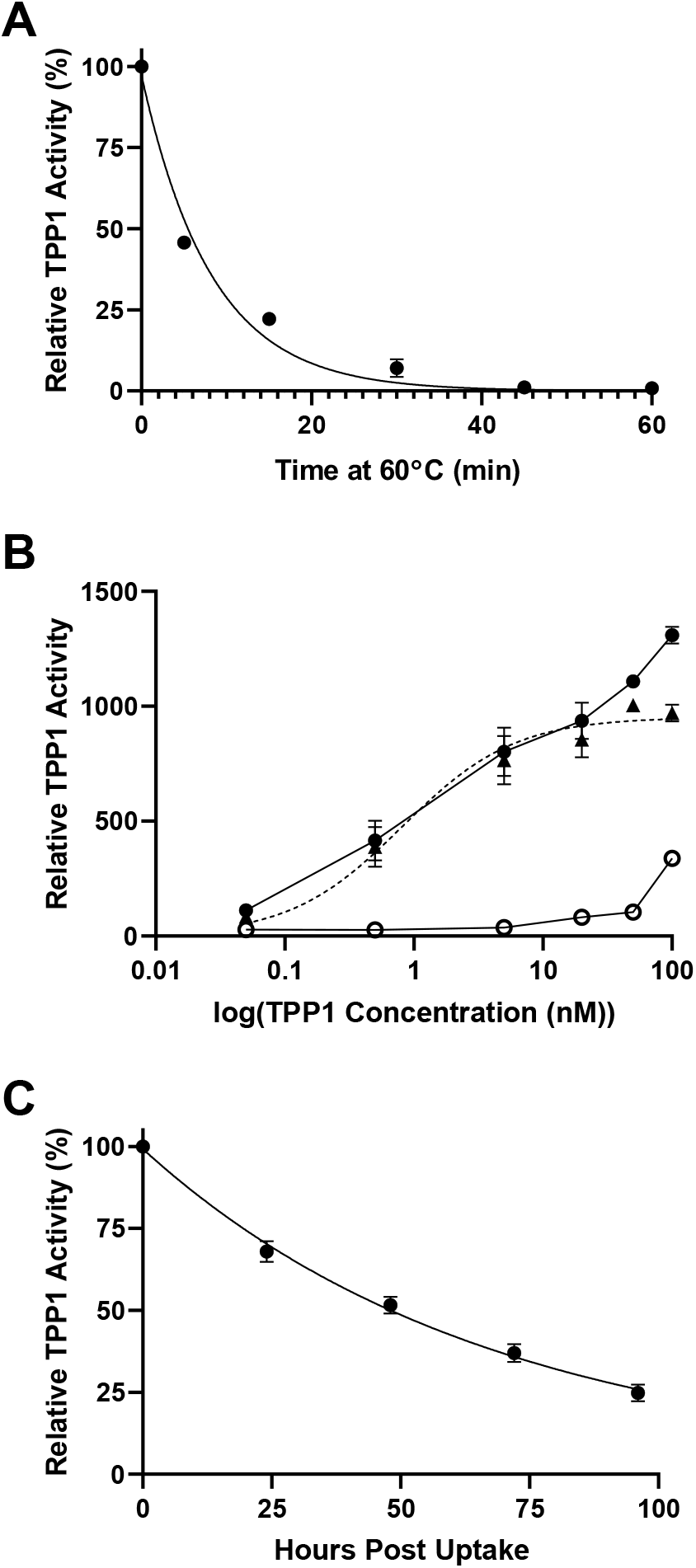
*In vitro* and lysosomal stability of WT TPP1. **(A)** Stability of enzymatic activity of mature WT TPP1 at 60°C. Data are compiled from two independent experiments conducted in duplicate. Duplicate measurements were averaged to generate a final decay curve. **(B)** Uptake of recombinant TPP1 by LINCL lymphoblasts. Cells were cultured with indicated concentration of proTPP1 for two days in the presence (open circle) or absence (closed circle) of 10 mM M6P before TPP1 activity was measured in cell extracts. Non-specific uptake was subtracted from total uptake in the absence of M6P to determine uptake specific to M6P receptor mediated endocytosis (closed triangle with dotted line). M6P specific uptake was fitted in Prism using a one-site specific binding model. Data are compiled from two independent experiments conducted in duplicate. **(C)** Activity of WT TPP1 in the lysosome after endocytosis by LINCL lymphoblasts. Data are compiled from two independent experiments conducted in duplicate. Symbols and error bars represent mean and standard deviation respectively in all cases.

### Reduced steric hindrance for β-turn structures

We designed and characterized TPP1 variants that are intended to optimize β-turn sequences to alleviate steric hindrance and improve conformational stability (Table 1). STRIDE ^27^ was first used to characterize secondary structural elements in proTPP1 based on its atomic coordinates ^7^. The location and type (as defined by the *ϕ* and *Ψ* angles of the second and third residues) of each turn structure was established and the amino acid composition of the β-turns analyzed ^28^. Amino acid frequency statistics for more than 7,000 β-turns from 426 crystal structures (26), indicate statistical preference for given amino acids that presumably stabilize the conformation of the turn. For each amino acid, a positional potential was calculated based on its frequency at a specific position within different β-turn types relative to its prevalence within the protein as a whole. As a strategy to potentially increase protein stability, amino acid residues within β-turns can be substituted with others that have a higher positional potential. This approach has been successfully applied to improve the thermal stability of bacterial and yeast proteins such as RNAse S_a_, S_a2_, S_a3_, ubiquitin, cold shock protein B, and histidine-containing phosphocarrier protein ^29,30^. When selecting amino acid changes to stabilize β-turn structures in TPP1, residues located in the interior of the protein or involved in hydrogen bonding were not considered. In addition, candidates for mutation to proline were not considered if the amide nitrogen of the native residue was a hydrogen bond donor.

**Table 1.**
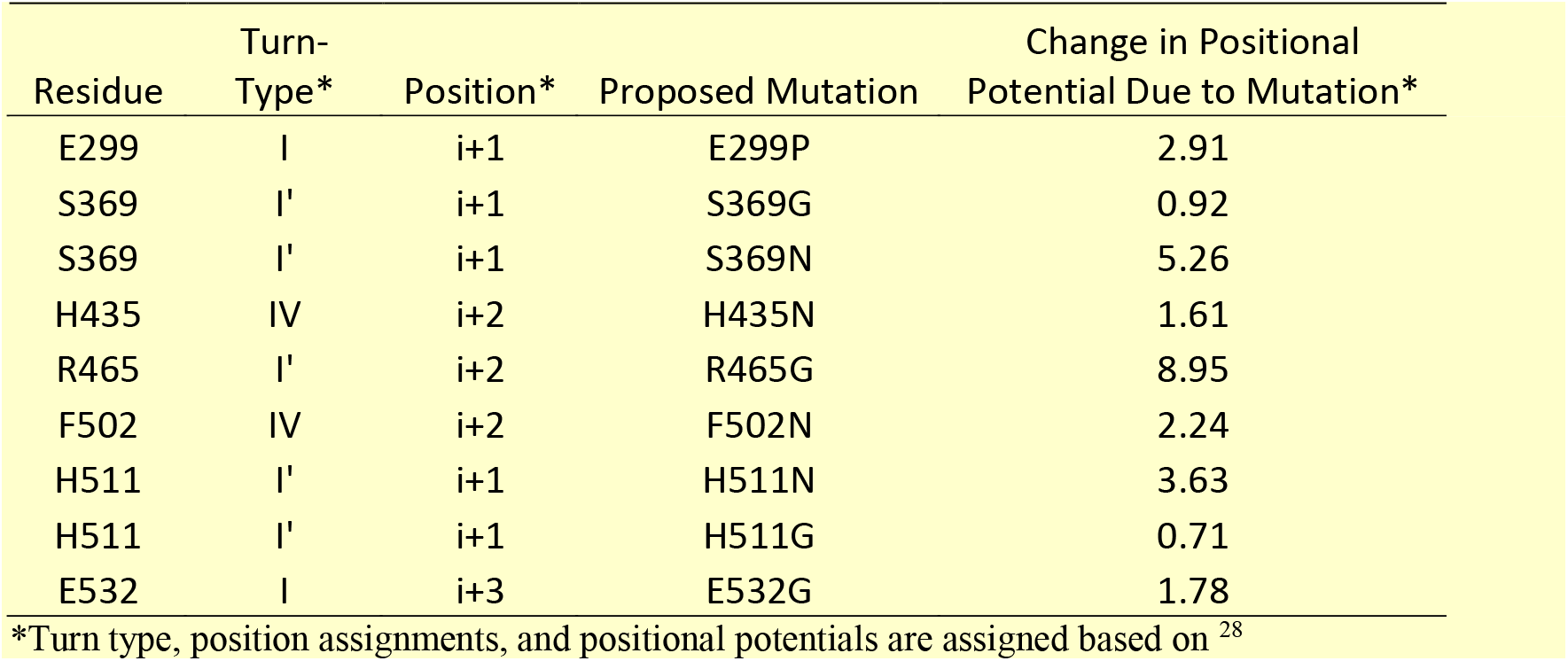
Effects of select mutations on positional potemntials.

We also examined the backbone conformation of the proTPP1 structure for residues with unusual rotational conformers where the dihedral angle *ϕ* > 0° (Figure S1). Such backbone conformations are infrequent in proteins, and usually are adopted by glycine as this amino acid lacks a sidechain and therefore has a greater range of permissible dihedral angles (Figure 2A) ^31,32^. Mutating these positions to glycine has previously been shown to improve the thermostability of bacterial proteins formate dehydrogenase and chondroitinase ABC 1 ^33,34^. Active site residues and asparagine residues identified as glycosylation sites were not considered. Identification of S369, R465, and H511 as candidate sites for mutation was consistent with the analysis above ^35^.

**Figure 2.**
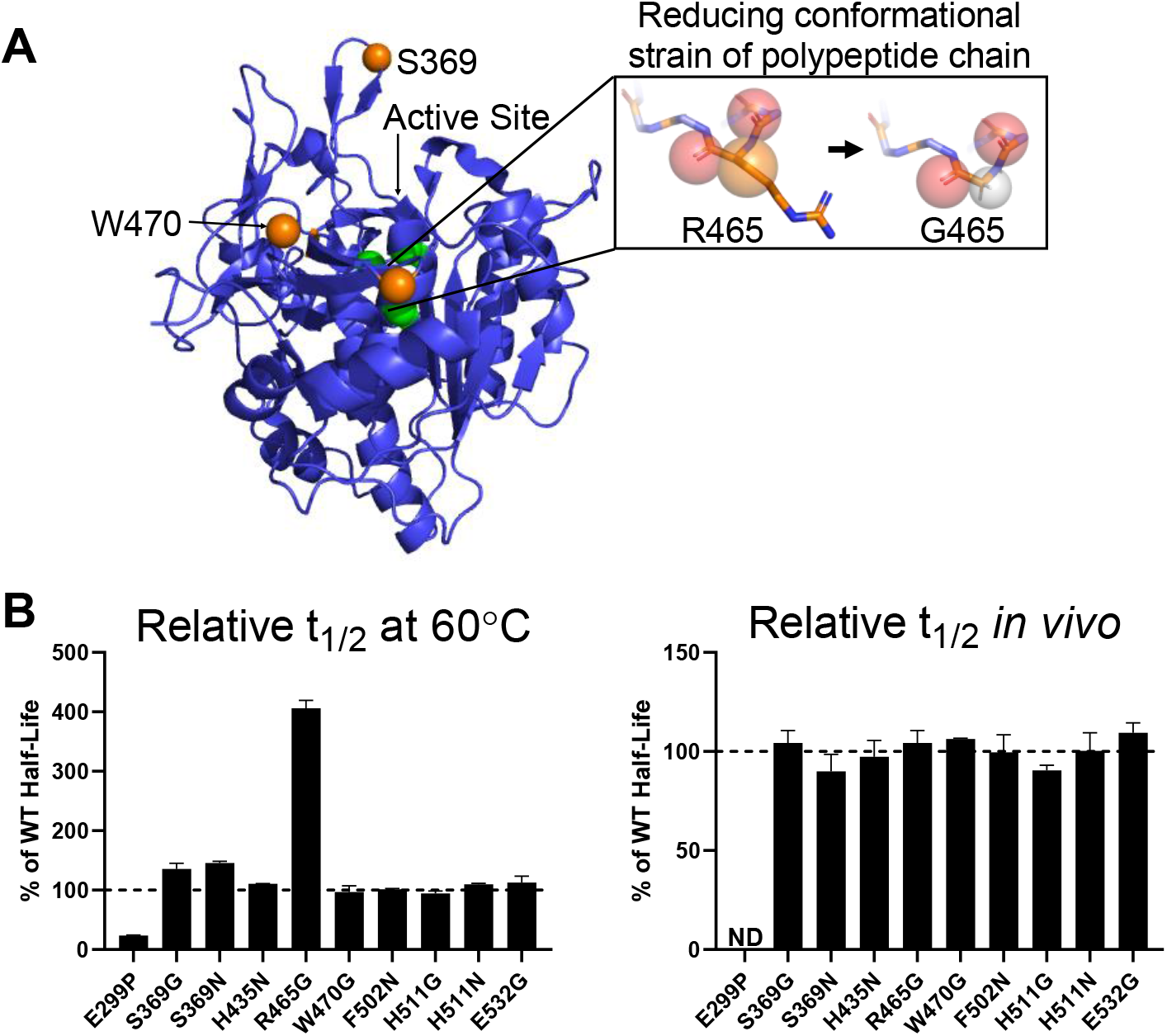
*In vitro* and lysosomal stability of the TPP1 β-turn stabilization variants. **(A)** Schematic of the methodology used to stabilize TPP1 utilizing the R465G mutation as an example. This is labeled on the crystal structure of proTPP1, rendered in Pymol, with the pro-domain removed, the active site labeled in green, and key β-turn stabilization variants labeled in orange. The spheres within the boxed area represent: red - the carbonyls for residues 464 and 465, orange - the β-carbon for R465, grey – the proton for G465. **(B)** Enzymatic half-life *in vitro* at 60°C and in the lysosome of LINCL lymphoblasts. ND = Not Determined

All 10 of the resulting variants were expressed at sufficient levels for characterization (Table S1). In *in vitro* thermal stability assays, six variants had half-life at 60°C that were similar to the 5.4 min half-life of WT TPP1, ranging between 5.0 and 6.1 min (Figure 2B). The mutation E299P substantially destabilized TPP1 and resulted in a half-life of only 1.2 min. Variants S369G and S369N showed slightly increased half-life of 7.7 and 7.5 min, respectively. The mutation R465G substantially improved the stability at 60°C with a half-life of 21.9 min. For the lysosomal half-life experiments, all TPP1 variants were similar to WT TPP1, with a half-life ranging between ~38 and ~52 hrs with the exception of E299P which displayed no activity following the initial two days of uptake (Figure 2C). Note that this included variant R465G, which displayed a significantly improved thermostability compared to WT TPP1 *in vitro*.

### Introduction of additional internal disulfide bonds

WT TPP1 contains three internal disulfide bridges (C111-C122, C365-C526, and C522-C537) ^7,8^. Previous studies have shown that introducing additional disulfide bonds with spatially proximal cysteine residues improved the thermostability of other proteins including T4 lysozyme, ribonuclease A, β-lactamase, lipase B, and insulin ^36–40^. There is substantial evidence that the increase in stability is due to a decrease in the entropy of the unfolded state of the protein, creating a greater free energy change between the folded and unfolded states of the protein. The program *Disulfide by Design 2.0* ^41^ was used to identify residue pairs which could potentially form additional disulfide bridges if converted to cysteine (Figure 3A). Residue pairs were selected that were located in loop regions of TPP1 and whose orientations and intermolecular distances were compatible with potential disulfide bond formation, without interfering with endogenous disulfide bonds. The 29 double cysteine variants selected for expression are listed in Table S1 but of these, only eight were expressed at sufficient levels for characterization.

**Figure 3.**
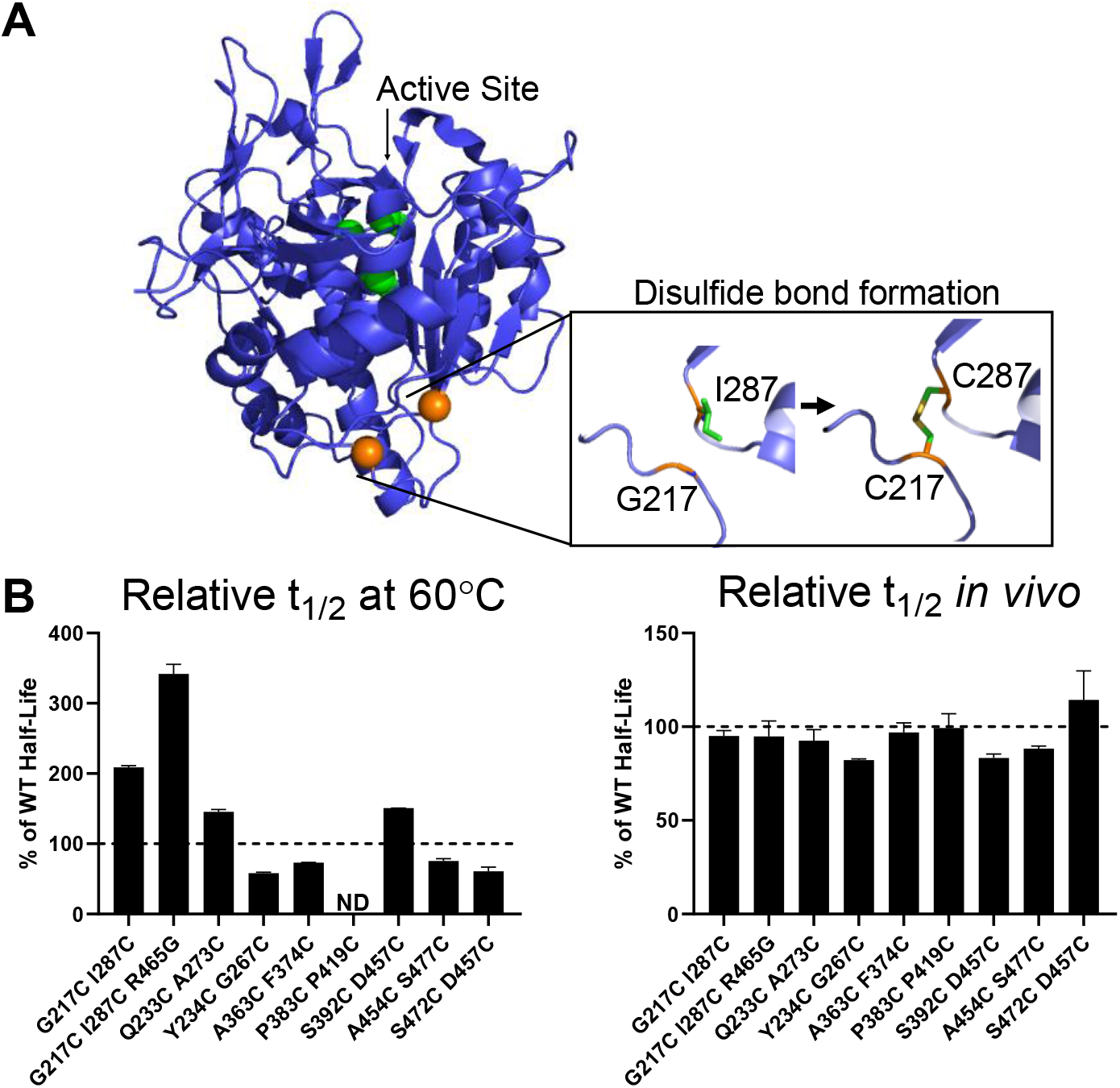
*In vitro* and lysosomal stability of TPP1 variants designed to introduce additional internal disulfide bonds. **(A)** Schematic of the methodology used to stabilize TPP1 utilizing the C217 I287 double variant as an example. Structure as in Figure 2A with the 217 and 287 residues labeled in orange. **(B)** Enzymatic half-life *in vitro* at 60°C and in the lysosome of LINCL lymphoblasts. ND = Not Determined

Most of the double cysteine variants tested had *in vitro* half-lives that were similar to WT TPP1, ranging between 3.0 and 4.3 min (Figure 3B). However, the Q233C A273C and S392C D457C variants had increased half-life *in vitro* of 8.1 and 7.8 min respectively, while G217C I287C had an *in vitro* half-life of 11.9 min. The G217C I287C mutations were then combined with R465G (see above) to determine if there would be a synergistic benefit but the *in vitro* half-life of the G217C I287C R465G triple variant was 18.6 min, which is similar to that of the R465G variant alone. All TPP1 double cysteine variants had lysosomal half-life that were similar to WT TPP1, ranging between ~30 and ~52 hrs (Figure 3C).

### Increasing structural homology with the thermoacidophilic peptidase Kumamolisin

Kumamolisin is a endopeptidase that, along with TPP1, is a member of the S53 sedolisin family ^6^. Kumamolisin was originally identified in the culture filtrate of the thermophilic bacterium *Bacillus novosp. MN-32* and displays optimal proteolytic activity at pH 3.0 and 70°C ^42,43^. Given that kumamolisin is thermostable, we reasoned that selective conversion of TPP1 sequences to resemble kumamolisin might increase TPP1 stability. Using the crystal structure of kumamolisin (PDB ID: 1GT9) ^44^ as a guide, we identified proline residues involved in facilitating turns that did not have corresponding proline residues in similar turns in TPP1. The location of the TPP1 residues that structurally aligned with these prolines in the Ramachandran plot was then used to identify which of these residues would favor proline substitutions. Conversely, while proline may have stabilizing properties in sharp turns of the polypeptide chain, it can destabilize in other contexts by disrupting α-helical turns. Kumamolisin has an isoleucine (I349) in place of a corresponding proline residue (P554) in a α-helical turn of TPP1. We therefore hypothesized that mutating this proline into an isoleucine might increase stability. These methodologies led to the creation of six variants (Table S1). Of these, three were active and showed similar half-life to WT TPP1 *in vitro* (Figure 4A), and in the lysosome (Figure 4B). The variant P554I showed a slightly decreased *in vitro* half-life of 4.20 minutes. This was likely due to P554 being located at the beginning of the α-helix where it played a role in stabilizing the structure.

**Figure 4.**
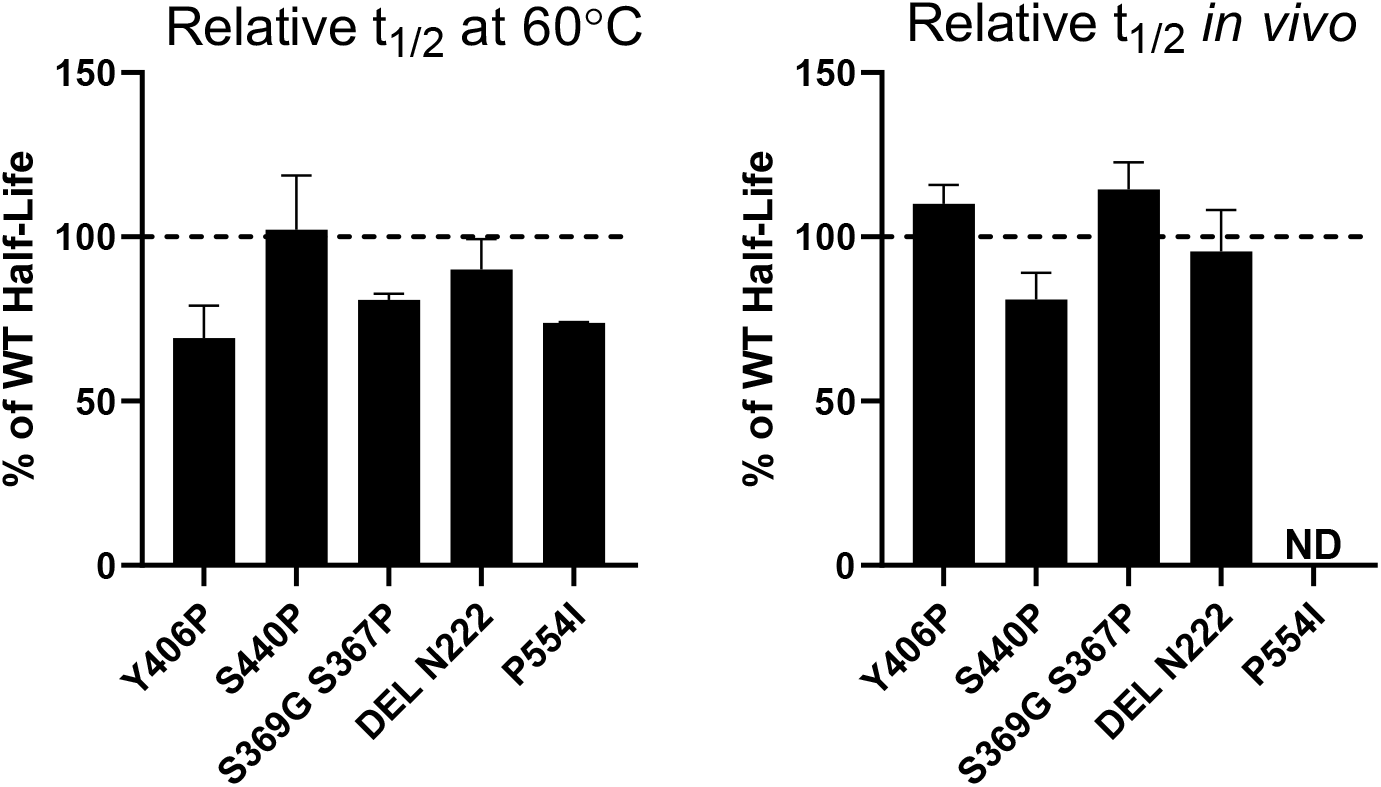
*In vitro* and lysosomal stability of the TPP1 variants designed to induce greater structural homology with kumamolisin. ND = Not Determined

Thermophilic proteins have been noted as often having shorter loop structures than their mesophilic counterparts ^45,46^. Based upon structural alignment between TPP1 and kumamolisin, loops were identified that were longer in TPP1 than in kumamolisin. Deletion mutations were designed to shorten these loops to resemble their counterparts in kumamolisin. We synthesized three TPP1 variants (Table S1) of which only one was active. This variant (Del N222) had a similar *in vitro* (Figure 4A) and lysosomal (Figure 4B) half-life as WT TPP1.

### Mutations to improve hydrophobic packing of the protein core

The program *Rosetta Void Identification and Packing* (VIP) is part of the RosettaDesign framework and assesses the core packing of a protein structure and proposes stabilizing mutations ^47^. A well-packed protein core can create complementary steric interactions and increase the buried hydrophobic surface area thus stabilizing the folded state of the protein (Figure 5A) ^48^. Using VIP, we identified eight mutations (Table S1) that would theoretically help to create stronger core packing in TPP1, six of which were expressed in sufficient quantities for characterization. Four variants displayed *in vitro* half-life similar to WT TPP1 while the M244F variant displayed a slightly increased *in vitro* half-life of 7.8 min (Figure 5B). All variants had a lysosomal half-life similar to WT TPP1 (Figure 5C).

**Figure 5.**
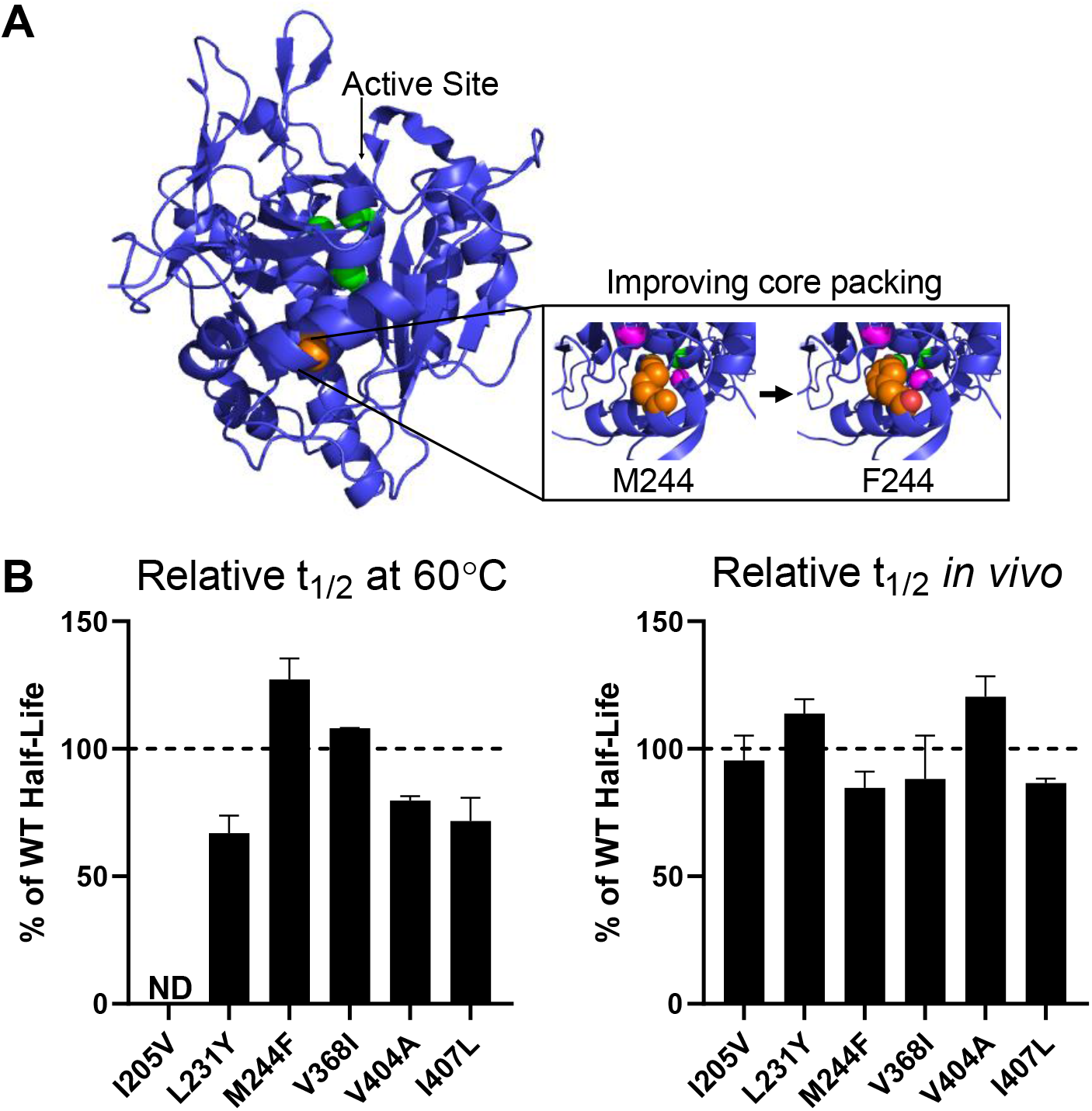
*In vitro* and lysosomal stability of the TPP1 variants designed to improve hydrophobic packing of protein core. **(A)** Schematic of the methodology used to stabilize TPP1 via the M244F mutation. Structure as in Figure 2A with the M244 residue labeled in orange. The cavities within 6Å of M244 are labeled in magenta. **(B)** Enzymatic half-life *in vitro* at 60°C and in the lysosome of LINCL lymphoblasts. ND = Not Determined

### Addition of a novel metal-binding loop

TPP1 has an octahedrally coordinated calcium-binding site that is conserved throughout the S53 sedolisin family ^7,8^. For other serine proteases, calcium binding commonly plays a role in facilitating endopeptidase activity and can also function to stabilize the protein ^49–52^. Engineering additional metal-binding sites has previously proven effective for improving the stability of proteins such as iso-1-cytochrome c, α-amylase, and collagen ^53–55^. Using the zinc-binding site of a kumamolisin variant specifically engineered to bind zinc as opposed to calcium (PDB ID: 4NE7) ^56^ as a guide, we inserted the amino acid sequence PAT between D457 and G458 in TPP1 to create part of a metal-binding loop. For the second loop, we added the amino acid sequence NDGPDGG between positions D405 and S408 of TPP1 and introduced mutations Y406N and I407A. Together, these changes were predicted to create an additional metal-binding site within TPP1. However, this variant was poorly expressed and insufficient amounts were obtained for characterization.

### Introducing additional salt bridges

Thermophilic proteins frequently have a greater number of salt bridges which can stabilize the protein by reducing the heat capacity change of unfolding ^57,58^. Introducing new salt bridges has been shown to increase the thermal stability of ribosomal protein L30e, nitrile hydratase, and β-glucosidase ^57,59,60^. We surveyed the surface region of TPP1 for residues that might exhibit high flexibility and low thermal stability ^60^. We analyzed the same regions of kumamolisin and incorporated salt-bridges found in kumamolisin into TPP1 by mutating corresponding residues. Of the two double variants (Table S1), only one (S392R Y406D) was expressed at sufficient levels for characterization, but its half-life *in vitro* and within the lysosome were the same as WT TPP1 (Table S1).

### Introducing “stabilizing” dipeptides

In an analysis of the dipeptide composition of 12 unstable and 32 stable proteins, certain dipeptide combinations were correlated with the overall stability of the protein ^61^. From this data, dipeptide instability weight values could be calculated for each dipeptide combination. The study noted that methionine, glutamine, proline, glutamate, and serine residues are found with a higher frequency in unstable proteins whereas asparagine, lysine, and glycine residues occur more frequently in stable proteins ^61^. Based upon dipeptide instability weight values, the primary sequence of TPP1 was examined for “destabilizing” peptides which were then mutated to generate “stabilizing” combinations with the highest possible weight value. Of the seven variants generated (Table S1), four expressed sufficiently for characterization, with three having similar half-life to WT TPP1 *in vitro* (Figure 6A) and within the lysosome (Figure 6B). Variant S525W had a reduced half-life both *in vitro* (2.0 min) and in the lysosome (~23 hrs).

**Figure 6.**
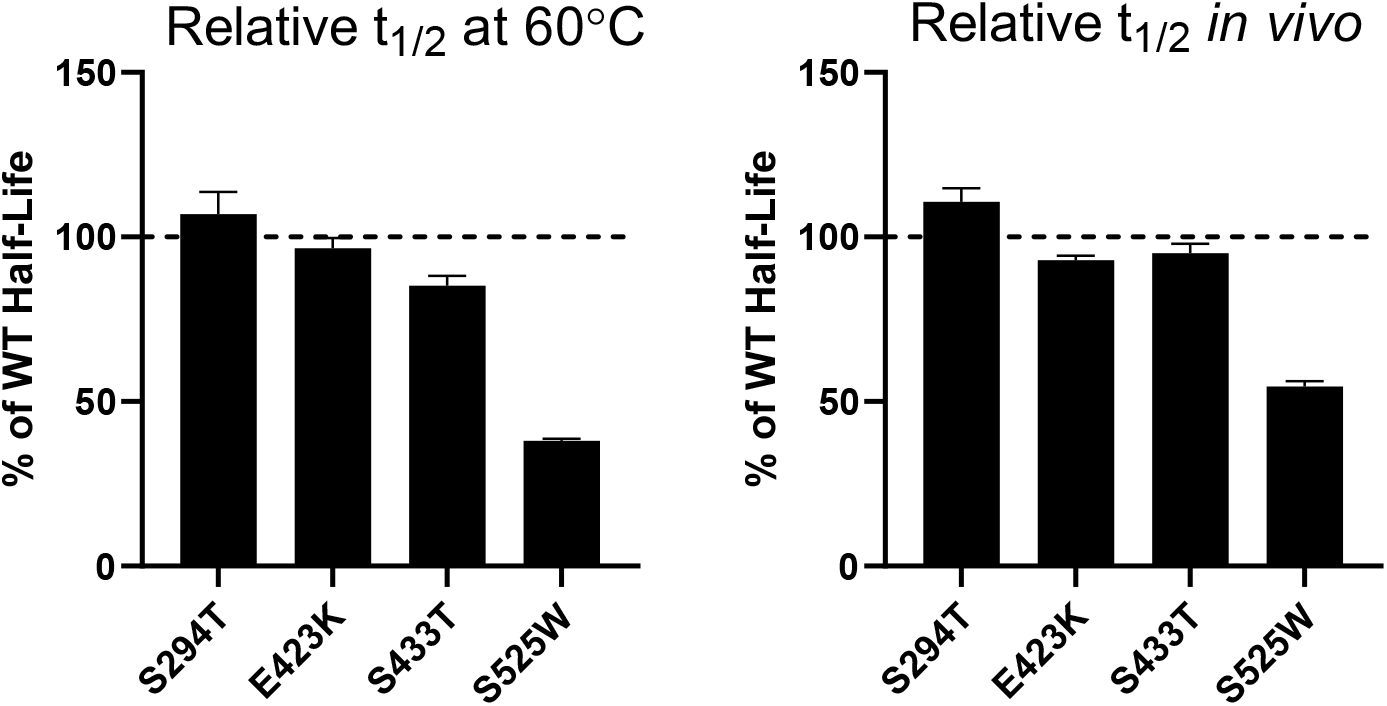
*In vitro* and lysosomal stability of the TPP1 variants designed to introduce “stabilizing dipeptides” into the primary structure.

### Adding potential glycosylation sites to improve stability

TPP1 contains N-linked glycans at surface asparagine residues 210, 222, 286, 313, and 443 which have been implicated in the maturation, activity, targeting, and stability of the protein ^7,8,14,62,63^. Previous studies have shown that introducing additional glycans by the insertion of novel canonical N-linked glycosylation sites (NXS/T) can increase the thermostability of proteins such as α-glucosidase, β-lactoglobulin, peanut peroxidase, and IgG1-Fc ^64–67^. Glycosylation may increase protein thermostability by increasing structural rigidity and compactness which stabilizes the folded state of the protein ^68,69^. Glycans may also protect proteins from proteolytic degradation ^70^, and this could be particularly important in the degradative environment of the lysosome.

To engineer new potential glycosylation sites into TPP1, we identified surface-exposed asparagine residues which are not already present in the context of a canonical N-linked glycosylation sequence. Using the NetNGlyc 1.0 Server ^71^, which predicts possible *N*-glycosylation sites, we introduced proximal mutations to create the NXS/T motif at these surface asparagine residues. *In vitro* (Figure 7A) and cellular (Figure 7B) half-life of the three resulting variants (Table S1) were similar to or shorter than that of WT TPP1.

**Figure 7.**
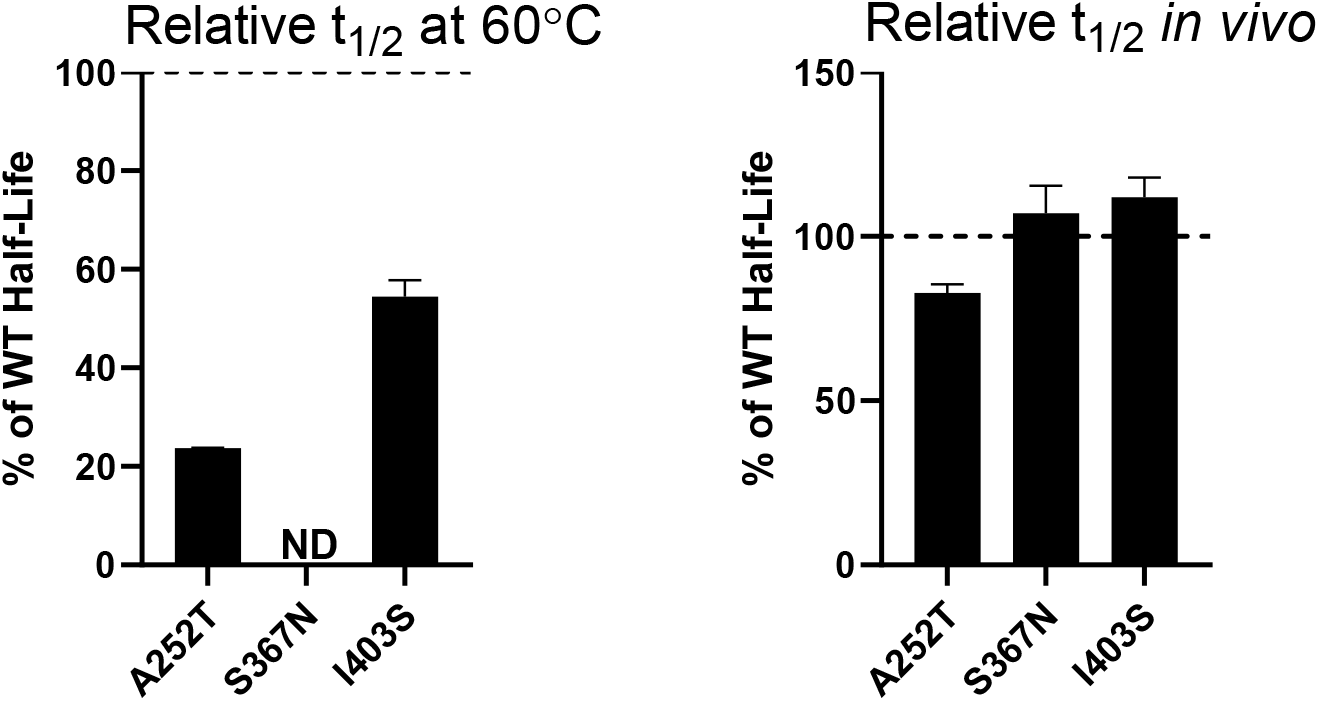
*In vitro* and lysosomal stability of the TPP1 variants designed to introduce additional glycosylation sites. ND = Not Determined

### Effects of stabilizing mutations on TPP1 thermal stability

Our protein engineering efforts identified a number of TPP1 variants with an increased *in vitro* half-life at 60°C. To examine the effects of the corresponding mutations on thermostability as measured by CD, WT TPP1 and TPP1 variants were expressed with a C-terminal hexahistadine tag and purified. TPP1 variants were activated and CD experiments were conducted to measure the T_m_ while TPP1 activity assays were performed to monitor thermal stability during denaturation (Table 2). Given the increased *in vitro* half-life conferred by the R465G mutation, the other β-turn stabilization mutations were also examined to determine if they provided subtle increases in protein stability that might not be detectable when stability was measured as a function of activity. Thermal denaturation monitored by CD showed a T_m_ of 56.2°C for mature untagged TPP1 (Figure 8A). Denaturation of mature his-tagged WT TPP1 indicated a T_m_ of 55.6°C (Figure 8B) measured by CD and 55.9°C via enzyme assay indicating that this sequence had minimal or no effect on stability. Similar studies were conducted for a total of 13 TPP1 variants (Figure 8C). Based on both CD and activity assays, eight of the variants had T_m_ values ranging from 53.8°C to 56.2°C, which is similar to that of WT TPP1. One variant (E299P) was found to decrease the thermal stability of TPP1. The mutations R465G and W470G increased TPP1 thermal stability based on CD measurements. R465G also showed increased stability according to activity measurements but the W470G mutation had a T_m_ that was almost identical that of WT TPP1. This result is consistent with the *in vitro* half-life results obtained at 60°C where the R465G mutation increased the enzymatic half-life of TPP1 while the W470G mutation did not. It is possible that the R465G mutation stabilizes regions of TPP1 that are essential for enzymatic activity while the W470G mutation generally stabilizes the secondary structure of the protein (hence increased T_m_ by CD) but does not stabilize regions important for catalytic function (hence no effect on functional stability). Variants S392C D457C and G217C I287C R465G were also found to increase the T_m_ of TPP1 as measured by both CD and functional assay. These results are consistent with *in vitro* half-life data at 60°C and show that combining G217C and I287C does not offer a synergistic increase in stability when combined with R465G.

**Table 2.** Effects of TPP1 Mutations on Thermal Denaturation Temperature.

| Mutation | $T_m$ (Activity) | $T_m$ (Ellipticity) |
| --- | --- | --- |
| WT TPP1 | $55.9 \pm 0.05$ | $55.6 \pm 0.17$ |
| M244F | $55.9 \pm 0.01$ | $54.9 \pm 0.09$ |
| E299P | $50.8 \pm 0.24$ | $52.0 \pm 0.04$ |
| S369G | $56.3 \pm 0.14$ | $55.9 \pm 0.28$ |
| S369N | $56.0 \pm 0.15$ | $54.4 \pm 0.11$ |
| H435N | $55.9 \pm 0.24$ | $54.4 \pm 0.09$ |
| R465G | $59.7 \pm 0.06$ | $64.4 \pm 0.04$ |
| W470G | $56.5 \pm 0.12$ | $63.4 \pm 0.11$ |
| F502N | $55.8 \pm 0.16$ | ND |
| H511G | $55.7 \pm 0.22$ | $55.7 \pm 0.14$ |
| H511N | $55.5 \pm 0.20$ | $54.9 \pm 0.44$ |
| E532G | $55.7 \pm 0.21$ | $53.8 \pm 0.09$ |
| S392C/ D457C | $56.7 \pm 0.15$ | $60.5 \pm 0.39$ |
| G217C/ I287C/ R465G | $59.3 \pm 0.23$ | $62.8 \pm 0.43$ |

**Figure 8.**
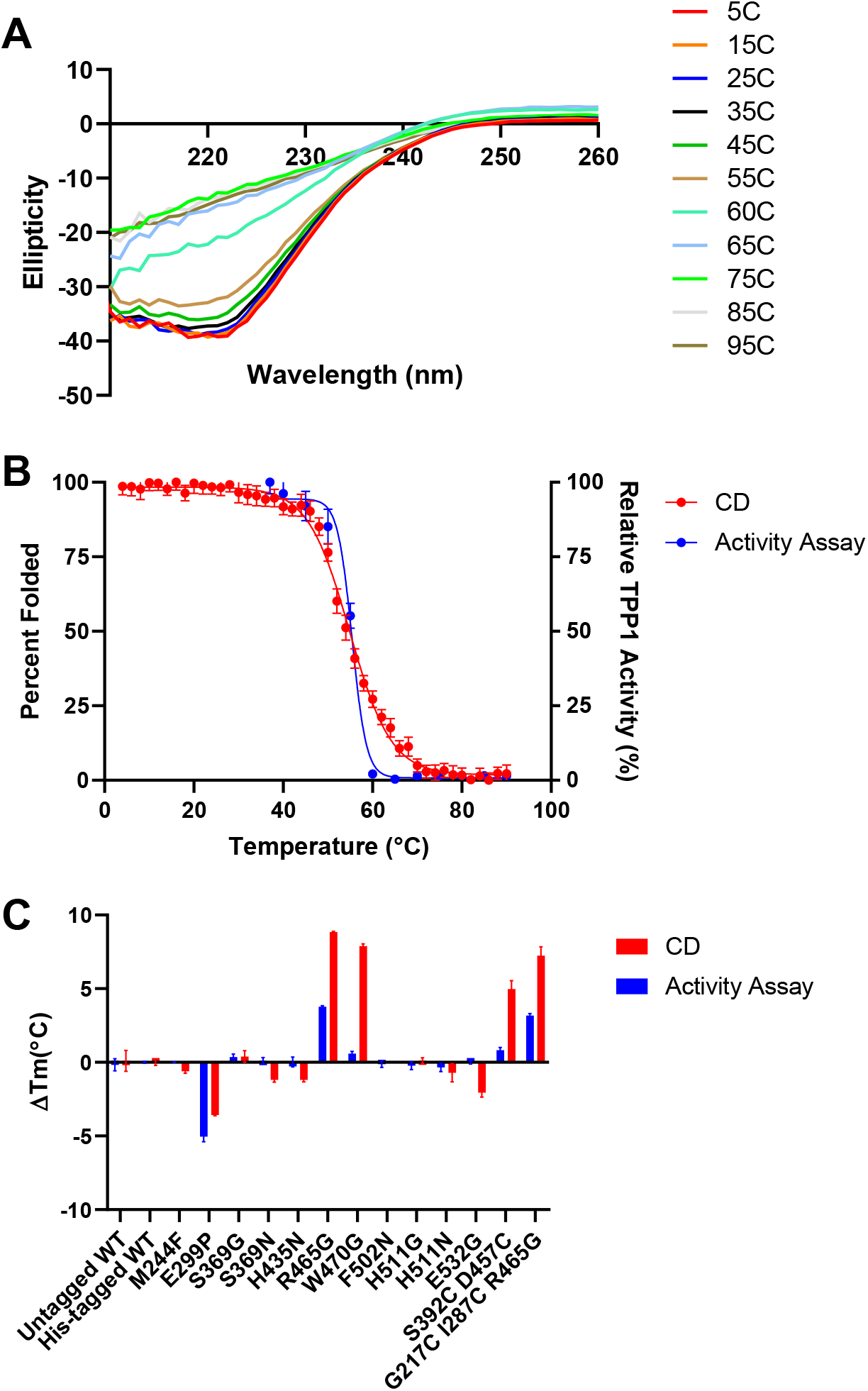
Thermal stability of mature, purified TPP1. **(A)** Untagged WT TPP1 CD spectra. Wavelength scans from 210-260 nm were performed on mature TPP1 at increasing temperatures and plotted against ellipticity. **(B)** Thermal denaturation of WT TPP1 determined by CD and enzymatic activity. Thermal denaturation experiments were carried out by monitoring the ellipticity of mature native TPP1 at 222 nm as the temperature was increased from 4°C to 90°C in 2°C increments. An equivalent sample of mature WT TPP1 was also heated from 40°C to 90°C in 5°C increments with TPP1 activity measured at each increment. **(C)** Change in melting temperature of his-tagged TPP1 variants compared to untagged WT TPP1. CD and enzyme activity assays where carried out for each TPP1 variant in the same manner as outlined in (B). Error bars display standard deviation in all cases. All thermal denaturation experiments were performed in triplicate.

## Discussion

Creating a TPP1 variant with improved pharmacokinetic properties could provide significant clinical benefits compared to the recombinant WT TPP1 that is currently used in the treatment of LINCL. For example, a TPP1 variant with an increased physiological half-life could potentially provide the basis for a more effective therapy, decreasing the amount of protein required per treatment, and/or an increasing the interval between doses, leading to an improved quality of life for LINCL patients. In this study, we utilized eight different protein engineering approaches to design over 70 TPP1 variants that were predicted to improve the thermostability of TPP1. Eight variants were identified that significantly increased the enzymatic half-life of TPP1 at 60°C, with R465G more than quadrupling the half-life. When variants of interest were purified and analyzed by CD, we found four variants that increased the T_m_ of TPP1. R465G provided the most significant effect by increasing the TPP1 T_m_ from 55.6°C to 64.4°C. Given its improved thermostability, the R465G variant may offer a more stable alternative to WT TPP1 with a longer shelf-life for those manufacturing and distributing the therapeutic protein.

Our *in vitro* screening process included evaluating the variants in terms of structural stability as determined by their resistance to thermal denaturation (CD assay) and functional stability as determined by their resistance to thermal inactivation (TPP1 activity assay). In general, structural and functional stability were well correlated but we identified one mutation (W470G) that stabilized the tertiary structure of TPP1 based on CD measurements but had no effect on its functional stability as measured by enzyme activity. W470 forms a packing interaction with Pro186 and Leu198 that spans the pro- and catalytic domains (Figure S2). The mutation of W470 to G may interfere with this interaction and may explain the behavior observed for the W470G variant. This suggests that the loss of enzymatic activity does not require global denaturation, but that local changes in secondary structure, likely near the active site or another essential functional element are enough to inhibit enzymatic activity.

We also measured the half-life of TPP1 variants in the lysosomal environment of LINCL lymphoblasts. Despite positive results from *in vitro* stability studies, none of the variants increased the lysosomal half-life of TPP1. It is interesting to note that amino acid changes that increased the thermostability of TPP1 *in vitro* were predicted using several different protein engineering methods, including stabilization of β-turn structures, introduction of additional disulfide bonds, and improved hydrophobic packing of the protein core, but none of these methods increased the physiological half-life of TPP1. This is likely because there are factors in addition to thermal stability that determine the functional life-span of TPP1 in the lysosome.

The lack of correlation between thermal stability of TPP1 *in vitro* and cellular half-life within the lysosome is intriguing and may reflect unique properties of this organelle. To determine whether this apparent disconnect is specific to lysosomal proteins, we compared protein thermal stability and half-life on an organelle–specific basis. Estimates of the melting point of proteins in the environment in which they function were obtained from cellular thermal protein profiling studies on three different cell lines ^72–74^ while protein half-lives were obtained from stable isotope labeling by amino acids in cell culture (SILAC) studies of protein turnover in five non-dividing cell types ^75^. We compared lysosomal proteins with proteins located in other organelles using two different classification schemes. First, we used the high stringency assignment of 3029 rat liver proteins ^76^ to assign eight major cellular locations. Proteins assigned to the lysosome included soluble luminal proteins such as TPP1 as well as integral and peripheral membrane proteins. In the second classification scheme, we restricted lysosomal proteins to 66 soluble proteins that are targeted to the lysosome by the M6P pathway ^77^.

Our meta-analysis of the thermal proteome profile of human hepatoma HepG2 cells ^72^, K562 leukemia cells ^74^, and HeLa cells in the M phase and G1S phase ^73^ consistently indicated that proteins residing in the lysosomal lumen (M6P-Lyso) had the highest average T_m_ (Figure 9, Table S2). Note that TPP1 was found in data sets for all four cell lines and had a Tm ranging from 52.9 to 53.9, slightly lower than mature TPP1 incubated *in vitro* at acidic pH (Table 2). Proteins of organelles other than the lysosome had similar T_m_s, with the exception of the nucleus which had the lowest average T_m_.

**Figure 9.**
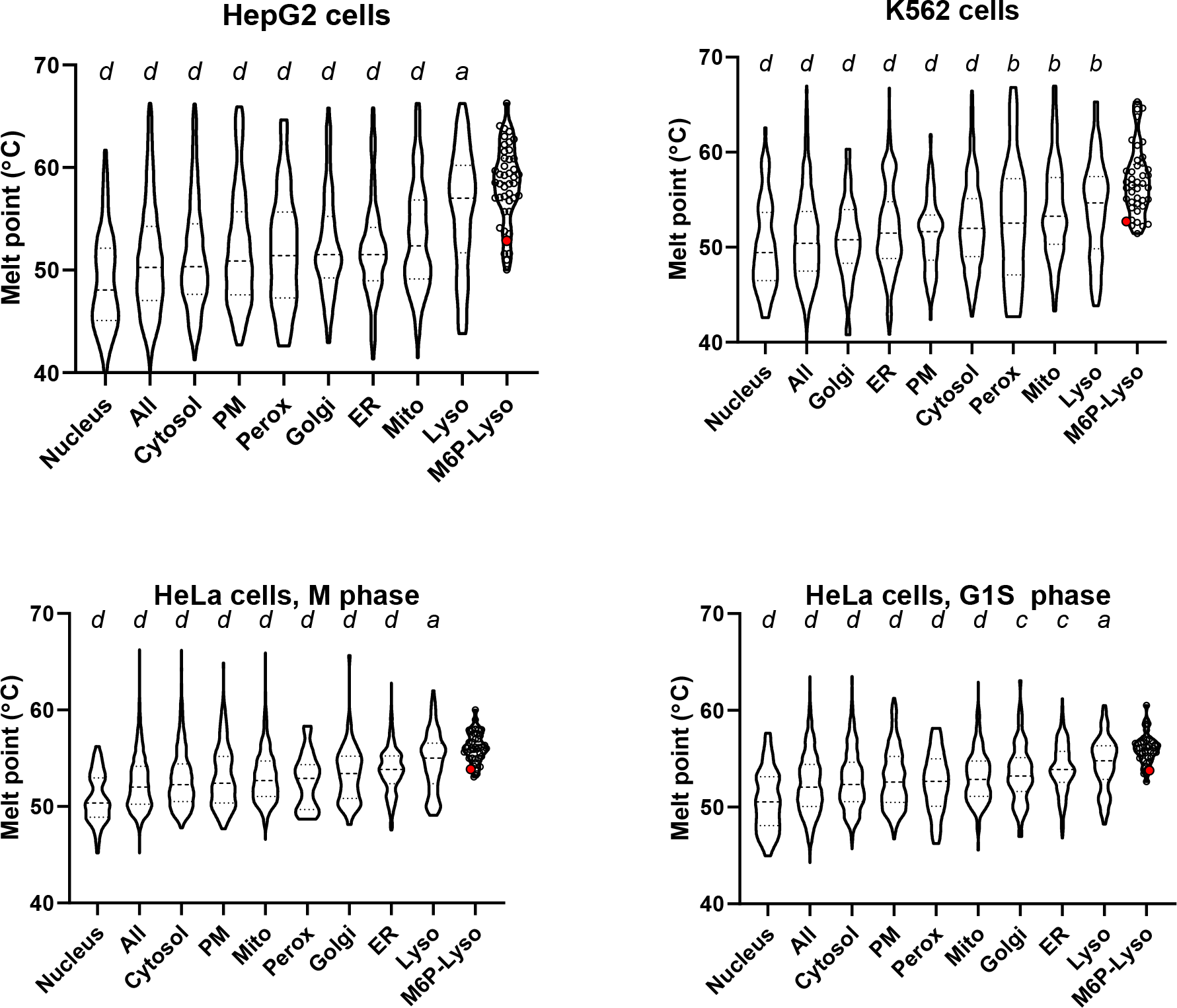
Meta-analysis of cellular thermal proteome profiling studies. Data used to generate plots are in Supplemental Spreadsheet 1. Quartiles are labeled via dashed lines. Values for individual proteins are shown by open circles for the M6P-Lyso set with the large red circle representing TPP1. Results using one-way ANOVA with Dunnett’s correction for multiple comparisons of M6P-lyso against all other categories are denoted above each violin plot: *a* ≤0.05; *b*≤ 0.01; *c*≤0.001; *d*≤0.0001. Abbreviations: PM, plasma membrane; Mito, mitochondria; Perox, peroxisomes; ER, endoplasmic reticulum; Lyso, lysosomes; M6P-Lyso, luminal lysosomal proteins targeted via the M6P pathway.

Analysis of protein turnover rates indicated that the average half-life of luminal lysosomal proteins compared to other types of proteins varies greatly with cell type (Figure 10, Table S2). In human B-cells and monocytes, the M6P-Lyso subset of proteins had the shortest lifetimes while they are intermediate in human hepatocytes and NK cells (Figure 10). In contrast, the M6P-Lyso proteins have the longest lifetimes in mouse neurons (Figure 10). Note also that the half-life values for TPP1 in these cells vary (hepatocytes, 97 hours; NK cells, 135 hours; B cells, 66 hours) but are not markedly different from the values for wild-type TPP1 taken by lymphoblasts in this study (median half-life, 43.5 h, range 30.2-81.6).

**Figure 10.**
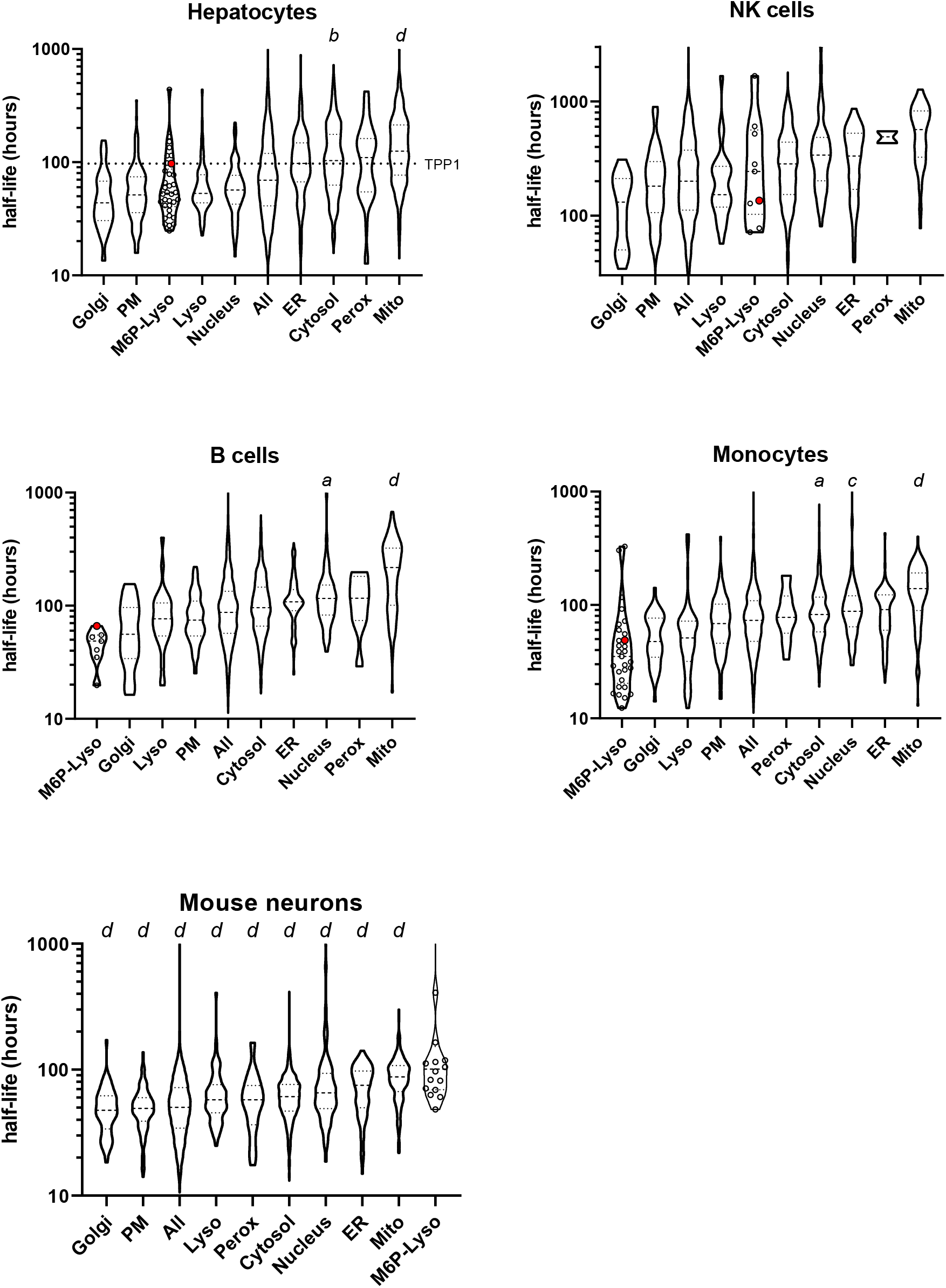
Meta-analysis of the protein half-life profile of non-dividing cells. Symbols as in Figure 9. Note that the TPP1 half-life was not reliably estimated in mouse neurons.

It is not clear why the half-lives of the luminal lysosomal proteins vary between the different cell types. Proteins of other organelles show less cell line to cell line variability. For example, Golgi proteins have the shortest half-life in all cell types while mitochondrial proteins have the longest, an observation also made by ^75^ when classifying their data set with GO terms for endoplasmic reticulum, Golgi apparatus, cytoplasm, mitochondrion and nucleus. It is possible that in neurons, the extended half-lives of M6P-Lyso proteins may relate to the fact that the M6P modification persists after delivery of newly synthesized lysosomal proteins to the lysosome ^78^ while in most other cell types, the M6P modification is removed by acid phosphatase 5, an enzyme that is low in brain ^79^. The retention of M6P may inhibit the action of glycosidases that further digest carbohydrate residues on lysosomal proteins whose presence may protect them from protease digestion ^70^. However, it is worth noting that when considering TPP1 specifically, the half-life of the enzyme in brain was similar to that of other tissues examined (brain, 72 hrs, liver, 79 hrs; heart, 60 hrs; kidney 108 hrs) ^80^. Taken together, this meta-analysis suggests that although lysosomal proteins are, on average, more thermally stable than the proteins of other organelles this does not result in an increased half-life and this is consistent with our observations with TPP1 in the current study.

It remains unclear why the functional stability of TPP1 variants measured in lymphoblasts does not reflect thermal stability measured *in vitro.* It is possible that there are region(s) of the protein that are critical for enzymatic function which are susceptible to protease digestion and are effectively a “weak point” for the protein. Thus, the mutations we have tested in this study, while stabilizing TPP1 as a whole, may have simply failed to address such a “weak point” that is limiting the ability of TPP1 to remain enzymatically active in the lysosome. Given that luminal lysosomal proteins have already evolved to have higher thermal stability than other classes of eukaryotic proteins, it is possible that there has been selective pressure to increase stability and promote survival in the protease-rich lysosome. It should also be emphasized that the majority of the protein engineering strategies utilized in this study to design the TPP1 variants were developed and validated using prokaryotic model proteins whose stabilities were measured by *in vitro* experiments. Such studies do not (and in most cases, do not need) to account for the complex physiological setting for proteins of interest, and this may be a significant factor for lysosomal proteins that function in an acidic degradative environment, which presents a unique challenge compared to other organelles. Future studies to understand how TPP1 is degraded within the lysosome would likely yield valuable insights into the regions of the protein that are the most susceptible to other lysosomal proteases and would therefore be the most beneficial to stabilize. Other orthogonal approaches, e.g., directed evolution, glycan engineering, or more in-depth structure analysis focusing on attributes other than thermal stability, may also assist in the discovery of more optimal TPP1 variants. In addition, conducting these higher-throughput primary screens under physiological conditions may also provide a more successful route to identifying modifications that impart the desired improved physiological properties.

## Acknowledgements

We would like to thank Dr. Sagar Khare and Dr. Manasi Pethe for providing technical assistance with regards to using Rosetta VIP for variant design.

## Conflict of Interest

P.L. and D.E.S. have received royalty payments as inventors on patent 8029781 (“Methods of Treating a Deficiency of Functional Tripeptidyl-Peptidase I [CLN2] Protein”), which is licensed to BioMarin Pharmaceutical Inc. The other authors declare no conflicts of interest.

## Funding

This work was supported by the National Institutes of Health [grant number NS037918 to P.L.]. The content is solely the responsibility of the authors and does not necessarily represent the official views of the National Institute of Health.

